# Microbial Peptidoglycan Engages Autophagy Receptor P62 to Induce Protective Mitophagy in the Liver

**DOI:** 10.1101/2025.06.27.662070

**Authors:** Jia Tie, Lang Wang, Xiong Wang, Meng Tian, Weiju Lu, Bin Qi, Zhao Shan

## Abstract

Although mitophagy is critical for maintaining mitochondrial integrity and hepatic homeostasis, the microbial-derived signals controlling this process remain unknown. Given the gut microbiota’s profound influence on liver pathophysiology, identifying specific bacterial factors that directly regulate hepatocyte mitophagy could unlock novel therapeutic strategies. In this study, we identify bacterial peptidoglycan (PGN)—a conserved cell wall component—as a key activator of mitophagy that protects against hepatocyte death. Through both *in vivo* and in *vitro* studies, we demonstrate that either heat-killed *Escherichia coli* or purified PGN attenuates hepatocyte death. Mechanistically, PGN is internalized by hepatocytes, localizes to mitochondria, and initiates mitophagy via direct interaction with the autophagy adaptor p62/SQSTM1. Genetic ablation of p62 in hepatocytes completely abolishes PGN-induced mitophagy, underscoring the pathway’s essential role. Strikingly, therapeutic administration of PGN markedly alleviates carbon tetrachloride (CCl4)-induced hepatic fibrosis, reducing collagen deposition and suppressing hepatic stellate cell activation through enhanced autophagic flux. Our work unveils a previously unrecognized host-microbe crosstalk in which PGN acts as a mitophagy inducer, offering a potential therapeutic avenue for liver diseases driven by mitochondrial dysfunction.

## Introduction

The liver’s unique anatomical connection to the intestinal tract through the portal circulation positions it as a primary target for gut-derived microbial signals(Hsu & Schnabl, 2023). Emerging research has established that the gut microbiota plays a crucial role in modulating hepatic physiology and protection against injury(Hu *et al*, 2025; Le *et al*, 2022; Yin *et al*, 2025). While this microbial influence is increasingly recognized, the specific bacterial components responsible for these hepatoprotective effects remain largely uncharacterized. This knowledge gap represents a critical limitation in our understanding of gut-liver crosstalk and its therapeutic potential.

Mitochondrial dysfunction is a hallmark of liver injury across diverse etiologies, including toxin exposure, metabolic overload, and carcinogenesis(Engelmann *et al*, 2021; Lin *et al*, 2014; Ma *et al*, 2024; Mansouri *et al*, 2018; Zhang *et al*, 2022). Damaged mitochondria accumulate reactive oxygen species (ROS) and pro-apoptotic factors like cytochrome c, which trigger cell death pathways(Bock & Tait, 2020). Additionally, impaired ATP production and mitochondrial dysfunction disrupt cellular homeostasis, further promoting hepatocyte injury and death(Schwabe & Luedde, 2018). Mitophagy, a selective form of autophagy that eliminates impaired mitochondria, is critical for mitochondrial quality control and cellular survival(Bock & Tait, 2020; Schwabe & Luedde, 2018). Pharmacological or genetic enhancement of mitophagy has shown therapeutic promise in alleviating liver injury, underscoring the need to identify endogenous or exogenous mitophagy inducers (Chen *et al*, 2024).

Gut microbial metabolites, such as short-chain fatty acids and secondary bile acids, are known to regulate mitochondrial energy metabolism and oxidative stress(Mann *et al*, 2024; Yang & Cong, 2021). For instance, butyrate enhances mitochondrial β-oxidation, while indole derivatives modulate mitochondrial membrane potential(Chimerel *et al*, 2013) and ATP production(Chimerel *et al*, 2014). Notably, recent studies revealed that bacterial peptidoglycan (PGN) fragments from *Escherichia coli* reduce mitochondrial stress, enhance ATP synthesis(Tian *et al*, 2024; Tian & Han, 2022), and promotes food digestion in *C. elegans*(Hao *et al*, 2024). However, whether microbial components directly regulate mitophagy—a process essential for mitochondrial health—remains unexplored. Unraveling this connection could provide new insights into the interplay between the gut microbiota and mitochondrial quality control, with significant implications for treating mitochondrial dysfunction-associated diseases.

We previously reported that heat-killed *E. coli* (HK *E. coli*) activates digestive system(Hao *et al*., 2024) and induces cellular stress response pathway(Liu *et al*, 2024) in *C .elegans,* enhancing the worms’ ability to adapt to environmental challenges. Building on these findings, we sought to investigate whether HK *E.coli* benefits mammals, particularly in mitigating liver injury. Surprisingly, we discovered that HK *E. coli* alleviates acetaminophen (APAP)-induced liver injury (AILI). Through both *in vitro* and *in vivo* studies, we further demonstrated that bacterial peptidoglycan (PGN), a major component of the bacterial cell wall, interacts with the autophagy receptor p62/SQSTM1 to trigger selective mitophagy, thereby preventing hepatocyte death. The data elucidated a novel mechanism by which gut microbial components directly influence hepatic cell survival via mitophagy, opening new avenues for microbiome-based therapeutic interventions in liver disease.

## Results

### PGN Attenuates Hepatocyte Death

To assess whether HK *E. coli* mitigate liver injury, mice were intraperitoneally injected with APAP and simultaneously orally gavaged with either PBS or HK *E. coli*. At 24 hours post-APAP administration, liver histology analysis revealed a significant reduction in necrotic area in HK *E. coli*-treated mice compared to PBS controls (Figure 1A). Consistent with this finding, serum ALT levels, a marker of liver injury, were significantly lower in HK *E. coli*-treated mice (Figure 1B). This suggests that HK *E. coli* prevents from AILI.

**Figure 1.**
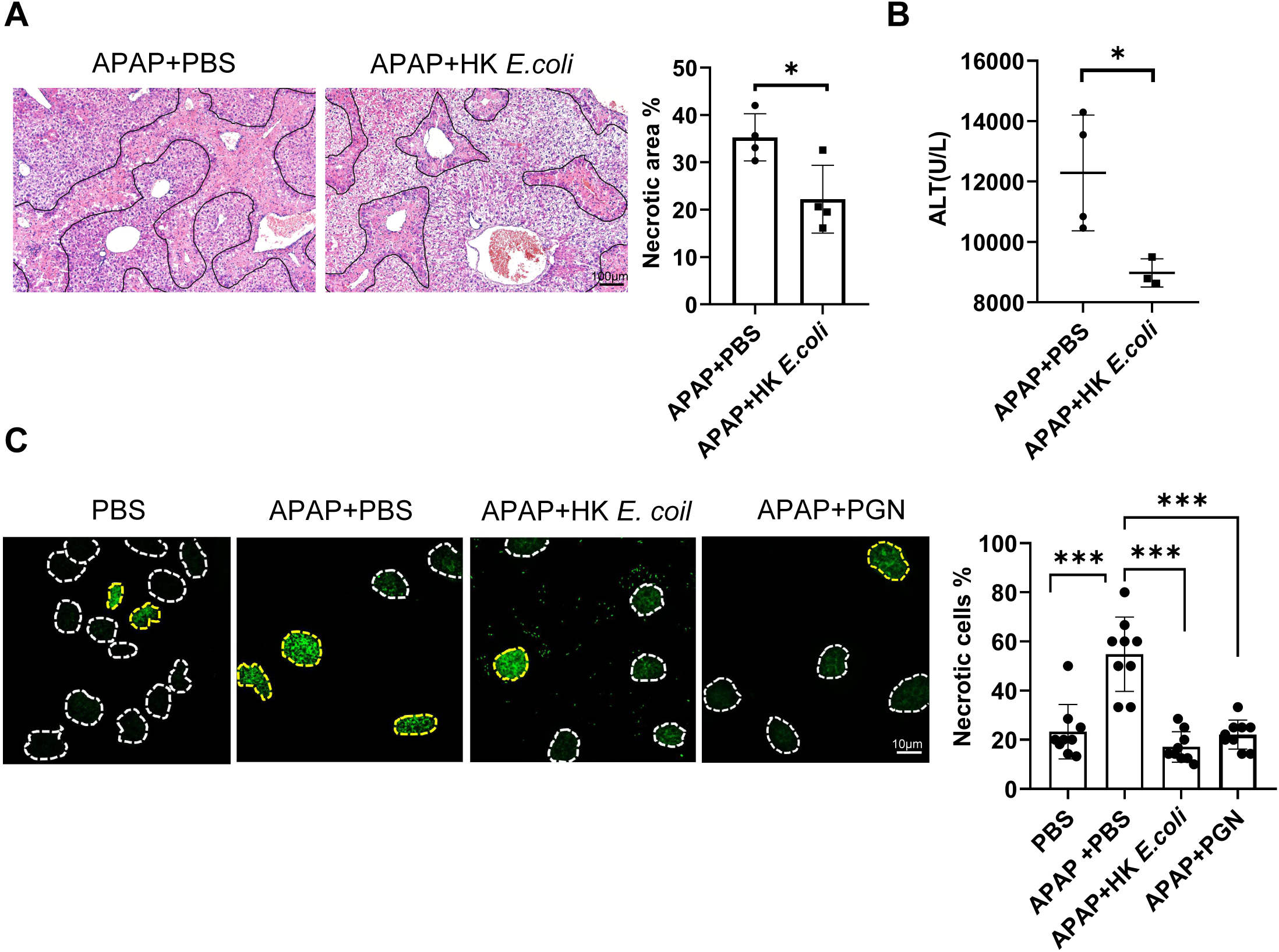
PGN attenuates hepatocyte death. **(A)** H&E staining of liver tissues from wild-type C57BL/6J mice treated with APAP and orally gavaged with PBS or heat-killed *E. coli* (HK *E. coli*) for 24 h. Quantification of necrotic area is shown (n = 4 mice/group). **(B)** Serum ALT levels in mice from (A). **(C)** SYTOX Green staining of Huh7 cells treated with PBS, APAP + PBS, APAP + HK *E.coli* or APAP + PGN. Necrotic cells (SYTOX Green+, yellow dashed outline) and viable cells (SYTOX Green−, white dashed outline) were quantified from 9 representative images/group. Data represent three independent experiments. Unpaired Student’s t-test (A, B); one-way ANOVA (C). *P < 0.05, ****P < 0.0001.

We next investigated the mechanism by which HK *E. coli* mitigates liver injury, considering two possibilities: modulation of APAP metabolism or direct effects on hepatocyte survival. APAP is metabolized by cytochrome P450 (CYP) enzymes into the toxic intermediate N-acetyl-p-benzoquinone imine (NAPQI), which is detoxified by glutathione (GSH). Excessive NAPQI accumulation leads to hepatocyte necrosis (Tujios & Fontana, 2011). To determine whether HK *E. coli* influences APAP metabolism, we analyzed key factors, including CYP2E1 (a major CYP enzyme involved in APAP metabolism), NAPQI-adduct formation, and GSH levels. Western blot analysis showed no differences in hepatic CYP2E1 protein levels (Figure S1A) or NAPQI-adduct formation (Figure S1B) between APAP + PBS and APAP + HK *E. coli*-treated mice at 2 hours post-APAP. Additionally, hepatic GSH levels remained comparable at all time points (Figure S1C). These results suggest that HK *E. coli* does not affect APAP bioactivation or detoxification.

To determine whether HK *E. coli* directly enhances hepatocyte survival, we assessed cell death using SYTOX Green staining, which labels necrotic cells. Compared to PBS-treated controls, HK *E. coli* significantly reduced the number of SYTOX Green-positive hepatocytes, indicating a protective effect against APAP-induced necrosis (Figure 1C). Since HK *E. coli* retains an intact cell wall rich in peptidoglycan (PGN)(Hao *et al*., 2024), we hypothesized that PGN might mediate this protection. Indeed, treatment with purified PGN alone mimicked the protective effect of HK *E. coli*, substantially reducing hepatocyte death (Figure 1C). These results demonstrate that PGN derived from HK *E. coli* mitigates APAP-induced hepatocyte necrosis independently of APAP metabolic pathways.

### PGN Targets Mitochondria and Interacts with Mitochondrial Proteins

To elucidate the mechanism by which PGN protects against hepatocyte death, we first examined its cellular uptake. Both in vitro hepatocyte treatment and in vivo oral gavage with FITC-conjugated PGN demonstrated efficient internalization, with robust fluorescence signal detected in hepatocytes (Figure 2A). This confirmed PGN’s ability to enter target cells. We then sought to identify PGN-interacting proteins by performing pull-down assays with liver tissue homogenates followed by LC-MS/MS proteomic analysis (Figure 2B). Among 658 identified PGN-associated proteins, a striking finding emerged: nearly 50% were mitochondrial-localized (Figure 2C). Gene Ontology (GO) enrichment analysis further supported this observation, revealing significant overrepresentation of mitochondrial-related processes including "cellular respiration" and "generation of precursor metabolites and energy," as well as mitochondrial compartments such as the "mitochondrial matrix" and "inner mitochondrial membrane protein complex" (Figure S2A). These results strongly suggest that PGN directly targets mitochondrial proteins and may modulate mitochondrial function, providing a potential mechanistic link to its protective effects against hepatocyte death.

**Figure 2.**
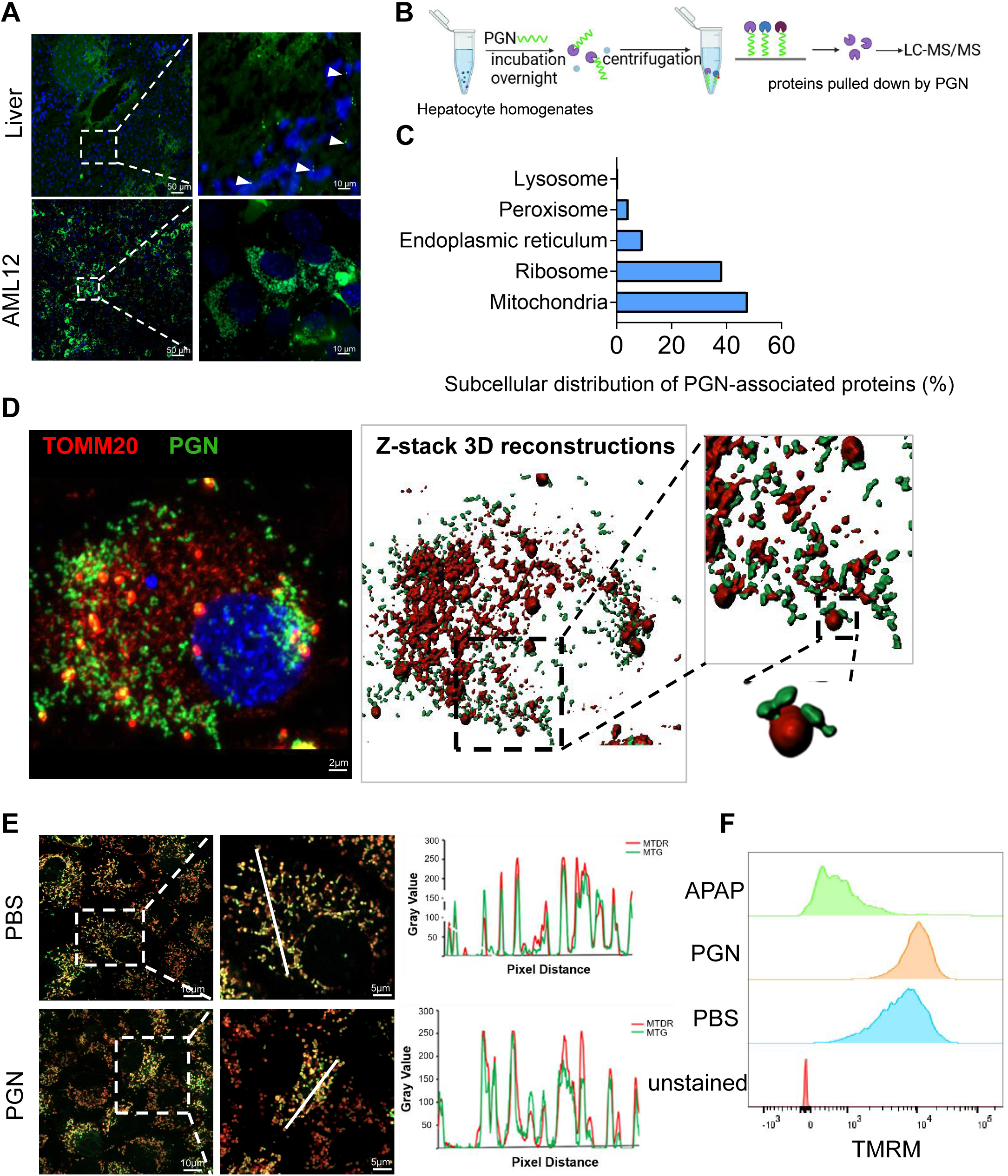
PGN targets mitochondria and interacts with mitochondrial proteins. **(A)** FITC-labeled PGN (green) was detected in mouse liver tissues and AML12 cells after 12 hours of oral gavage or direct treatment, with nuclei counterstained by DAPI (blue). **(B)** Experimental workflow for identifying PGN-interacting proteins by LC-MS/MS from primary hepatocytes. **(C)** Proteomic analysis showing subcellular distribution of PGN-associated proteins identified in (B). Full mass spectrometry data are available in Table 1. **(D)** Visualization of FITC-PGN (green) subcellular localization on mitochondria. TOMM20 (red) was used to label mitochondria outer membrane. Left panel: 2D fluorescence imaging; Middle panel: 3D reconstruction; Right: Magnified views with orthogonal projections (inset). Nuclei were counterstained with DAPI (blue). **(E)** Mitochondrial health were evaluated after 48-hour treatment with PGN or PBS control using (Left) MitoTracker Green (MTG, total mitochondria) and MitoTracker Deep Red (MTDR, membrane potential-dependent). (Right) Fluorescence intensity profiles along indicated transects (white lines). **(F)** Flow cytometry analysis of mitochondrial membrane potential using TMRM in Huh7 cells treated with APAP, PGN, or PBS (with unstained controls).

To confirm mitochondrial localization of PGN, we performed immunofluorescence staining with TOMM20, a mitochondrial outer membrane marker, in FITC-PGN-treated hepatocytes. Confocal microscopy demonstrated strong colocalization between FITC-PGN and TOMM20, which was further confirmed by 3D reconstruction and orthogonal projections (Figure 2D). To investigate potential effects on mitochondrial function, we employed dual-labeling with MitoTracker probes: Deep Red (specific to healthy mitochondria) and Green (labeling all mitochondria regardless of health status). Both PBS- and PGN-treated hepatocytes showed similar colocalization patterns, indicating that PGN treatment does not induce mitochondrial damage (Figure 2E). This observation was further supported by flow cytometry analysis of tetramethylrhodamine methyl ester (TMRM) fluorescence, which revealed that PGN treatment maintained mitochondrial membrane potential at levels comparable to PBS controls (Figure 2F). These findings collectively demonstrate that PGN specifically targets mitochondria and interacts with mitochondrial proteins while preserving mitochondrial integrity and function in hepatocytes.

### PGN Triggers Mitophagy in Hepatocytes

To investigate the effects of PGN on mitochondrial activity, we performed RNA-seq on hepatocytes treated with APAP + PBS or APAP + PGN (Figure S3A). Principal component analysis (PCA) revealed distinct transcriptional profiles between the two groups (Figure S3B). Consistent with our proteomics data showing PGN localization to mitochondria without inducing damage (Figure 2), KEGG pathway analysis demonstrated significant enrichment of the phagosome pathway among the top 20 upregulated pathways (Figure S3C). Given that (i) PGN physically associates with mitochondria (Figure 2B-D) and preserve mitochondrial integrity (Figure 2E-F) and (ii) phagosome pathway activation is closely linked to mitochondrial quality control(Minton, 2015), these results collectively suggest that PGN may promote mitophagy—a selective form of autophagy targeting mitochondria— without causing overt mitochondrial dysfunction.

To directly evaluate mitophagy induction, we employed Huh7 cells (a human hepatocyte cell line) stably expressing the pH-sensitive mitophagy reporter mt-Keima. This reporter distinguishes lysosomal (acidic) mitochondrial delivery via a shift in excitation peak (440 nm [neutral pH] → 586 nm [low pH])(Sun *et al*, 2024). Flow cytometry revealed a significant increase in the acidic/total mitochondria ratio upon PGN treatment, comparable to the carbonyl cyanide m-chlorophenyl hydrazone (CCCP) positive control (Figure 3A). Confocal microscopy further validated these findings, showing enhanced mt-Keima red fluorescence (pH 4) in PGN-treated cells, which reflects mitochondrial translocation to lysosomes (Figure 3B). To evaluate the completeness of PGN-induced mitophagy, we analyzed autophagic flux. Western blotting showed increased LC3-II (a marker of autophagosome formation) accompanied by a moderate elevation in p62 levels in PGN-treated cells (Figure 3C). When co-treated with bafilomycin A1 (BAF), a potent lysosomal inhibitor, we observed further accumulation of both LC3-II and p62, confirming that PGN stimulates complete mitophagic flux (Figure 3C). Complementary ultrastructural analysis by transmission electron microscopy (TEM) demonstrated a significantly higher frequency of mitophagosomes in PGN-treated cells compared to PBS controls (Figure 3D). This was further corroborated by immunofluorescence staining showing enhanced colocalization of mitochondria with lysosomal marker Lamp1 following PGN treatment (Figure 3E). Together, these data demonstrate that PGN activates mitophagy in hepatocytes, promoting efficient mitochondrial clearance.

**Figure 3.**
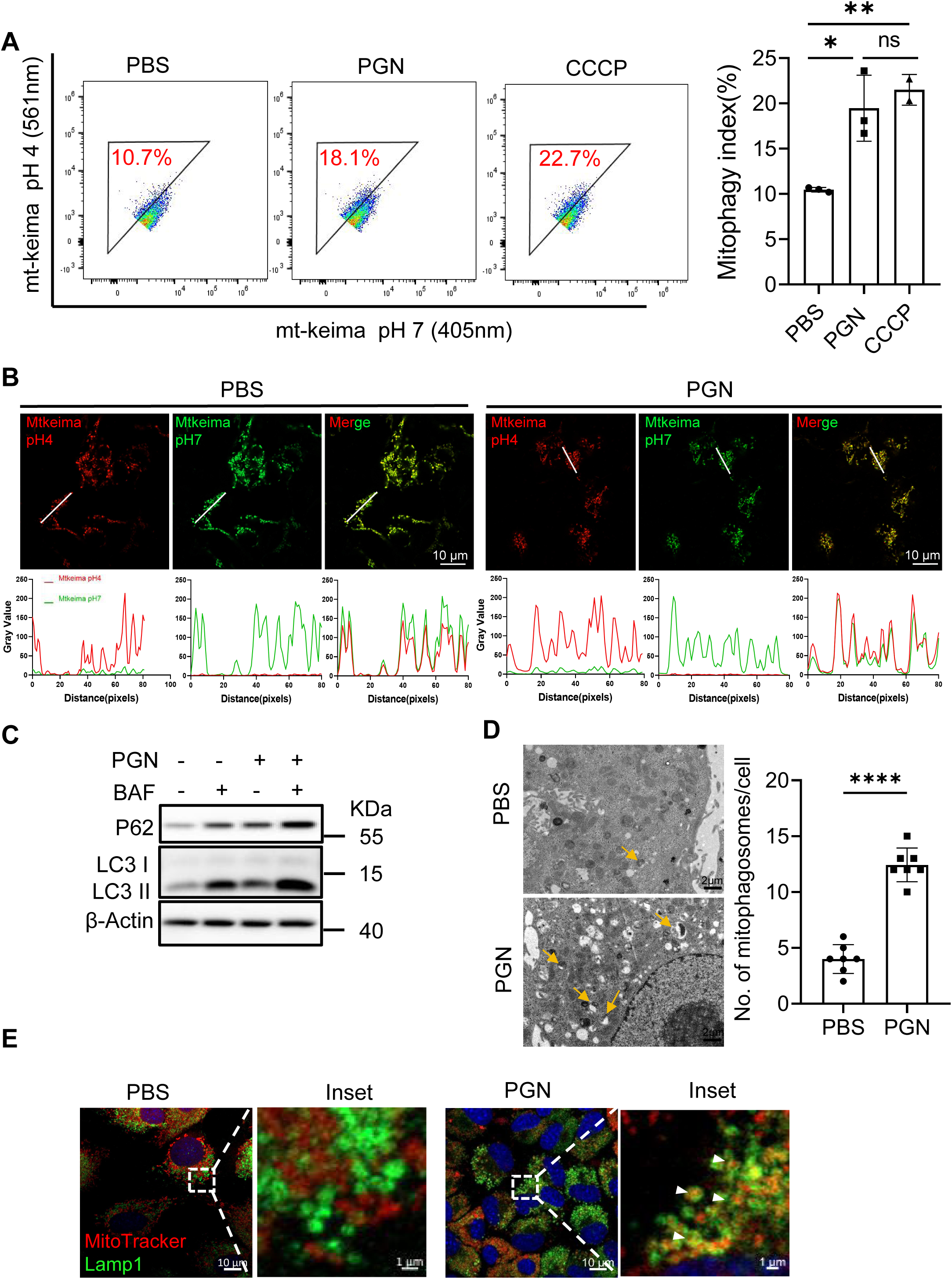
PGN triggers mitophagy in hepatocytes. **(A)** Flow cytometry to assess mitophagy induction by PGN in Huh7 cells using mt-Keima. Quantified mitophagy index (acidic/total mitochondria ratio) from three biological replicates. CCCP was used as positive control. **(B)** Representative fluorescence images to show mitophagy induction by PGN in Huh7 cells (Top). Fluorescence intensity profiles along indicated transects (white lines, bottom). **(C)** Western blot analysis of autophagy markers (P62, LC3-I/II) to assess autophagic flux in AML12 cells treated with PGN ± bafilomycin A1. **(D)** TEM visualization of mitophagosomes in AML12 cells treated with PBS or PGN. Left: Representative images (yellow arrowheads indicate mitophagosomes). Right: Quantification of mitophagosomes per cell section (n = 7 cells/group). **(E)** Mitochondrial-lysosomal colocalization in AML12 cells treated with PBS or PGN stained with MitoTracker (red) and Lamp1(green). White arrowheads indicate Mitochondrial-lysosomal colocalization. Unpaired Student’s t-test (D); One-way ANOVA (A). *P < 0.05, **P < 0.01, ****P < 0.0001; ns, not significant (P ≥ 0.05).

### PGN Directly Interacts with P62 to Promote Mitophagy

To elucidate the mechanism of PGN-induced mitophagy, we analyzed our proteomic data of PGN-interacting proteins and identified several autophagy-related factors, including the key autophagy receptor p62/SQSTM1 (Figure S4A). p62 is known to initiate autophagosome formation by recruiting LC3 to cargo targets(Palikaras *et al*, 2018). We systematically validated this interaction through complementary experimental approaches: First, endogenous co-immunoprecipitation assays in hepatocytes revealed specific binding between PGN and p62 (Figure 4A). To determine whether this interaction occurs directly, we performed *in vitro* binding assays using purified His-tagged p62 protein, which demonstrated strong and specific binding to PGN (Figure 4B). This direct physical interaction was further confirmed by fluorescence microscopy, where FITC-conjugated PGN showed clear colocalization with His-p62 immobilized on beads (Figure 4C). These results establish that PGN physically interacts with p62 through direct molecular binding, suggesting a potential mechanism for PGN-mediated regulation of selective autophagy.

**Figure 4.**
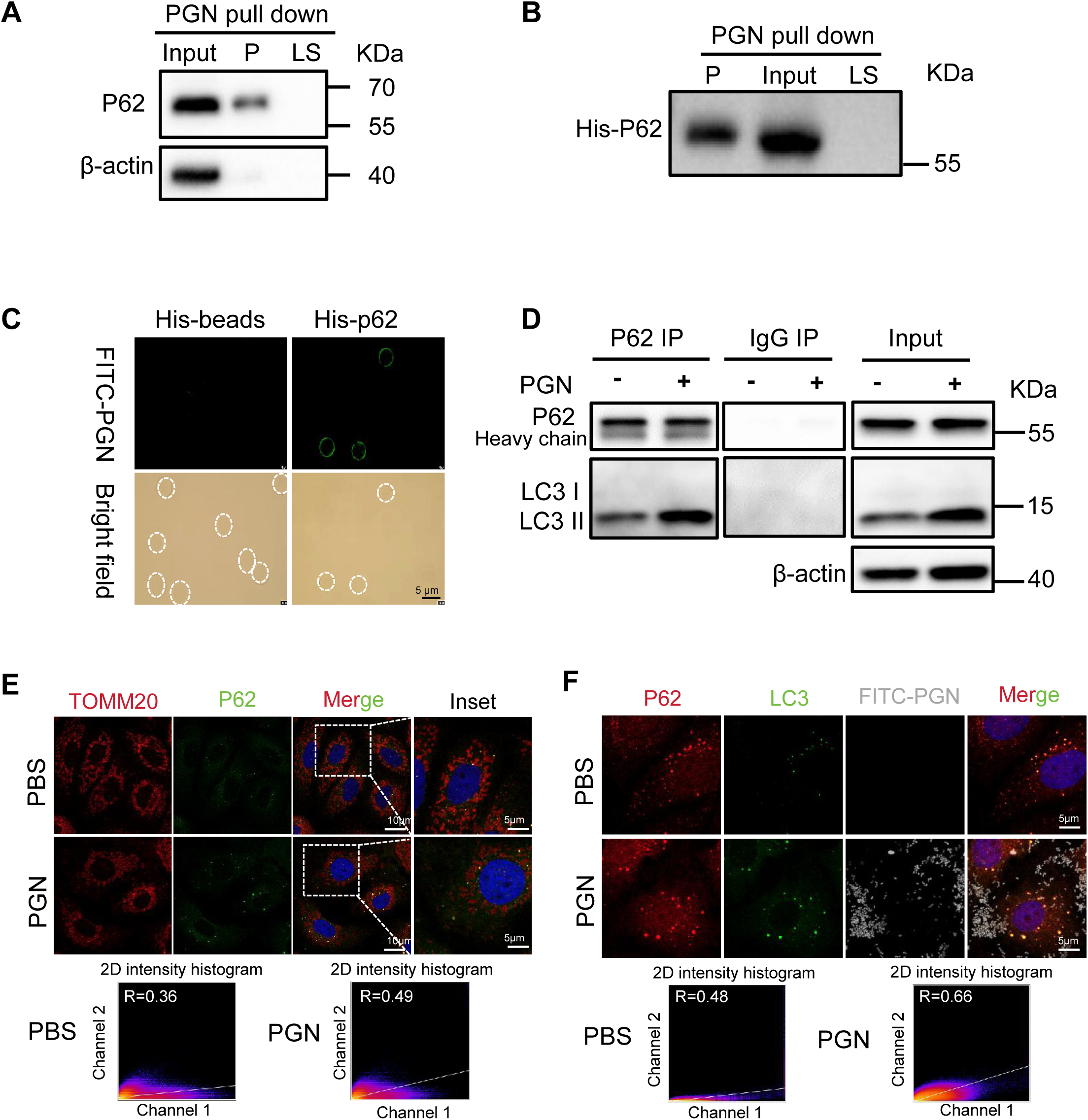
PGN directly interacts with P62 to promote mitophagy. **(A)** Western blot analysis of endogenous P62 pulled down by PGN in AML12 cells. P (pellet), LS (supernatant from last wash). **(B)** Western blot analysis of recombinant His-tagged P62 (His-P62) pulled down by PGN *in vitro*. P (pellet), LS (supernatant from last wash). **(C)** Microscopic validation of PGN-P62 interaction. FITC-labeled PGN bound to His-P62-conjugated beads was visualized by fluorescence microscopy. Beads were outlined with white dashed lines in bright-field images. His-tagged beads alone served as a control. **(D)** Western blot analysis of immunoprecipitated LC3 by anti-P62 antibody in AML12 cells treated with or without PGN. IgG was used as a negative control. **(E)** Representative fluorescence images to show P62 (green) recruitment onto mitochondria (TOMM20, red) by PGN in AML12 cells (Top). Colocalization was quantified using the Pearson correlation coefficient, with higher values indicating greater overlap. **(F)** Representative fluorescence images to show P62 (red) and LC3 (Green) colocalization under PGN treatment in AML12 cells (Top). Colocalization was quantified using the Pearson correlation coefficient. Data represent three independent experiments (A-F).

To functionally validate the PGN-p62 interaction in mitophagy induction, we performed a series of complementary experiments. First, co-immunoprecipitation analysis revealed that PGN treatment significantly enhanced the interaction between p62 and LC3, as evidenced by increased LC3-II recovery in p62 pulldowns (Figure 4D). This biochemical evidence was supported by elevated LC3-II conversion, indicating augmented autophagosome formation. We next employed high-resolution imaging to spatially characterize this process. Immunofluorescence analysis demonstrated that PGN treatment: (i) increased p62 recruitment to mitochondria (TOMM20+) (Figure 4E), and (ii) enhanced colocalization between p62 and LC3 at mitochondrial sites (Figure 4F). These findings visually corroborated the biochemical data, showing PGN’s ability to promote autophagosome assembly specifically at mitochondria.

### PGN-Induced Mitophagy Requires p62 *In Vivo*

To establish the physiological requirement of p62 in PGN-induced mitophagy, we developed a hepatocyte-specific *in vivo* monitoring system using AAV8 vectors expressing the mt-Keima mitophagy reporter under control of the hepatocyte-specific thyroxine-binding globulin (TBG) promoter (Figure 5A). Quantitative live imaging analysis demonstrated that oral PGN administration significantly increased the proportion of acidified mitochondria (red mt-Keima signal) compared to PBS-treated controls (Figures 5B, 5C), confirming PGN’s ability to induce hepatic mitophagy in living animals. To genetically validate p62’s essential role, we employed CRISPR-Cas9-mediated hepatocyte-specific p62 knockdown. Co-delivery of AAV-TBG-sgP62 and AAV-mtKeima to Cas9-expressing mice achieved approximately 70% reduction in p62 protein levels in primary hepatocytes (Figure 5D). Strikingly, while control mice (AAV-TBG-sgCtrl) exhibited robust PGN-induced mitophagy, this response was nearly completely abolished in p62-deficient livers (Figures 5E, 5F). These *in vivo* genetic studies provide definitive evidence that: (i) PGN potently induces hepatocyte mitophagy under physiological conditions. (ii) p62 is an essential mediator of this protective pathway.

**Figure 5.**
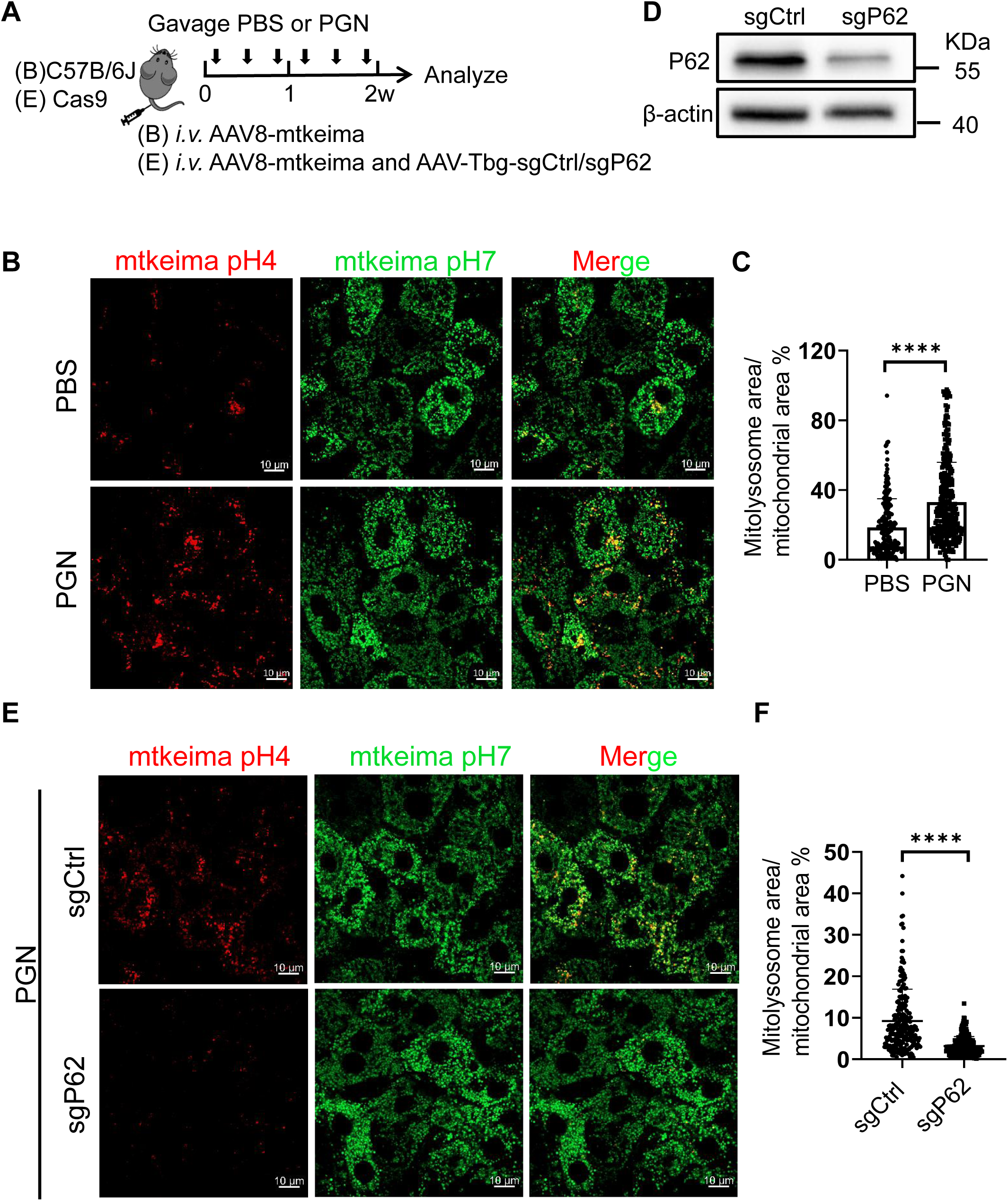
PGN-induced mitophagy requires P62 *in vivo*. **(A)** Experimental design for *in vivo* assessment of PGN-induced mitophagy and its dependence on P62. (B) WT model: C57BL/6J mice received intravenous AAV8-mtKeima to report mitophagy, followed by oral gavage with PBS (control) or PGN (3×/week for 2 weeks). (E) P62-KO model: Cas9 mice were co-injected with AAV8-mtKeima and AAV-Tbg-sgP62 (or AAV-Tbg-sgCtrl) to delete P62 in hepatocytes, then treated with PBS or PGN identically to WT mice. **(B)** Representative confocal live-cell images of hepatic mtKeima in PBS- or PGN-treated mice. **(C)** Quantification of hepatic mitophagy levels from (B). (n = 3 mice/group). **(D)** Western blot to detect P62 protein levels in primary hepatocytes from AAV-Tbg-sgCtrl or AAV-Tbg-sgP62-treated mice. **(E)** Representative confocal live-cell images of hepatic mtKeima in PGN-treated mice injected with AAV-Tbg-sgCtrl or AAV-Tbg-sgP62. **(F)** Quantification of hepatic mitophagy levels from (E). (n = 3 mice/group). Unpaired Student’s t-test (C, D). ****P < 0.0001.

### PGN Alleviates Liver Fibrosis

To determine whether PGN’s cytoprotective effects require functional mitophagy, we employed pharmacological inhibition using Mdivi-1, a selective inhibitor of mitochondrial fission that blocks mitophagy initiation. Western blot analysis revealed that Mdivi-1 treatment significantly attenuated PGN-induced mitophagy, as evidenced by: reduced LC3-II conversion (indicating impaired autophagosome formation) and accumulation of p62 (suggesting blocked autophagic flux) (Figure S5A). Consistent with these molecular findings, SYTOX Green viability assays demonstrated that while PGN treatment alone markedly reduced hepatocyte necrosis, concurrent Mdivi-1 administration completely abolished this protective effect (Figure S5B). These complementary biochemical and functional analyses establish that PGN’s anti-necrotic action strictly depends on intact mitophagy.

To assess the therapeutic potential of PGN in chronic liver injury, we employed a carbon tetrachloride (CCl4)-induced fibrosis model with PGN intervention initiated after two weeks of injury (Figure 6A). PGN administration significantly mitigated hepatocellular damage, reducing serum ALT activity by approximately 50% (Figure 6B). Histological analysis demonstrated that PGN administration markedly decreased collagen deposition, with Sirius Red staining quantifying a ∼35% reduction in fibrotic area (Figure 6C). Furthermore, immunohistochemical analysis demonstrated suppression of hepatic stellate cell (HSC) activation, evidenced by a ∼40% decrease in desmin-positive area (Figure 6D). Autophagy examination showed that while basal autophagy was activated during fibrogenesis (increased LC3-II and P62 accumulation), PGN treatment further enhanced autophagic flux, as indicated by elevated LC3-II coupled with reduced P62 levels (Figure 6E). These findings collectively establish that the PGN-p62-mitophagy axis as a therapeutically actionable pathway for liver diseases associated with mitochondrial dysfunction.

**Figure 6.**
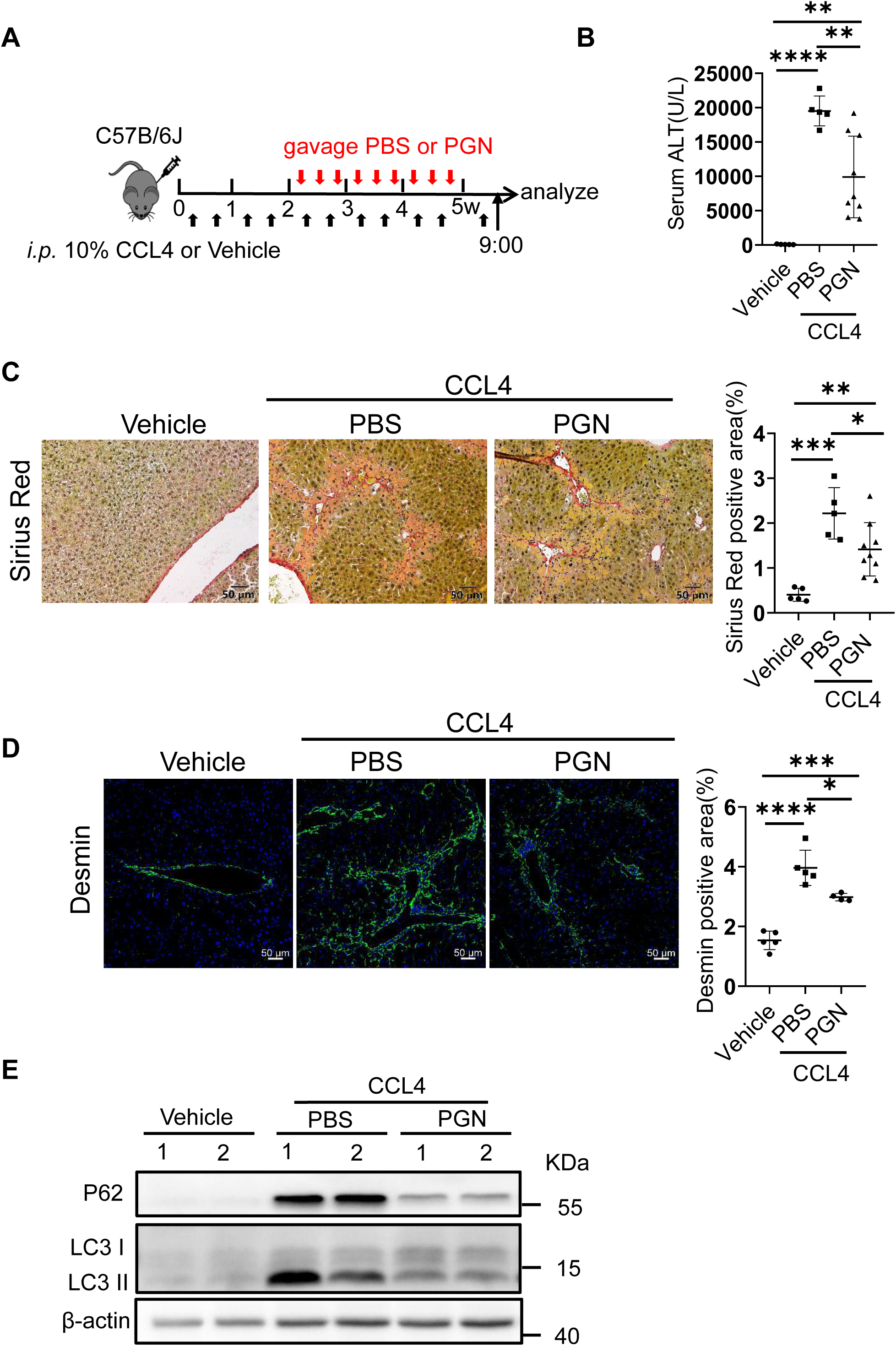
PGN alleviates liver fibrosis. **(A)** Experimental design for evaluating PGN’s therapeutic effect in CCL4-induced liver fibrosis. C57BL/6J WT mice received intraperitoneal (i.p.) injections of 10% CCl4 twice weekly for 11 doses. PBS or PGN was administered by oral gavage three times weekly from weeks 3 to 5 (*w*, week). **(B)** Serum ALT levels in mice from (A). n=5-9 mice/group. **(C)** Representative images to show liver collagen deposition assessed by Sirius Red staining. Sirius Red positive area was quantified. (n = 5-9 mice/group). **(D)** Representative images to show activated hepatic stellate cells (HSCs) by desmin immunofluorescence staining. Desmin-positive area was quantified. (n = 5 mice/group). **(E)** Western blot analysis of autophagy markers (P62, LC3-I/II) to assess autophagic flux in murine livers of mice treated in (A). One-way ANOVA (B-D). *P < 0.05, **P < 0.01, ***P < 0.001,****P < 0.0001.

## Discussion

The findings presented in this study reveal a previously unrecognized mechanism by which the gut microbiota-derived peptidoglycan (PGN) exerts hepatoprotective effects through selective activation of mitophagy. Our data demonstrate that PGN is internalized by hepatocytes, localizes to mitochondria, and initiates a p62/SQSTM1-dependent mitophagy that clears damaged organelles, thereby mitigating liver injury (Figure 7). This discovery not only expands the known biological functions of bacterial cell wall components but also provides new insights into host-microbe interactions in maintaining hepatic homeostasis.

**Figure 7.**
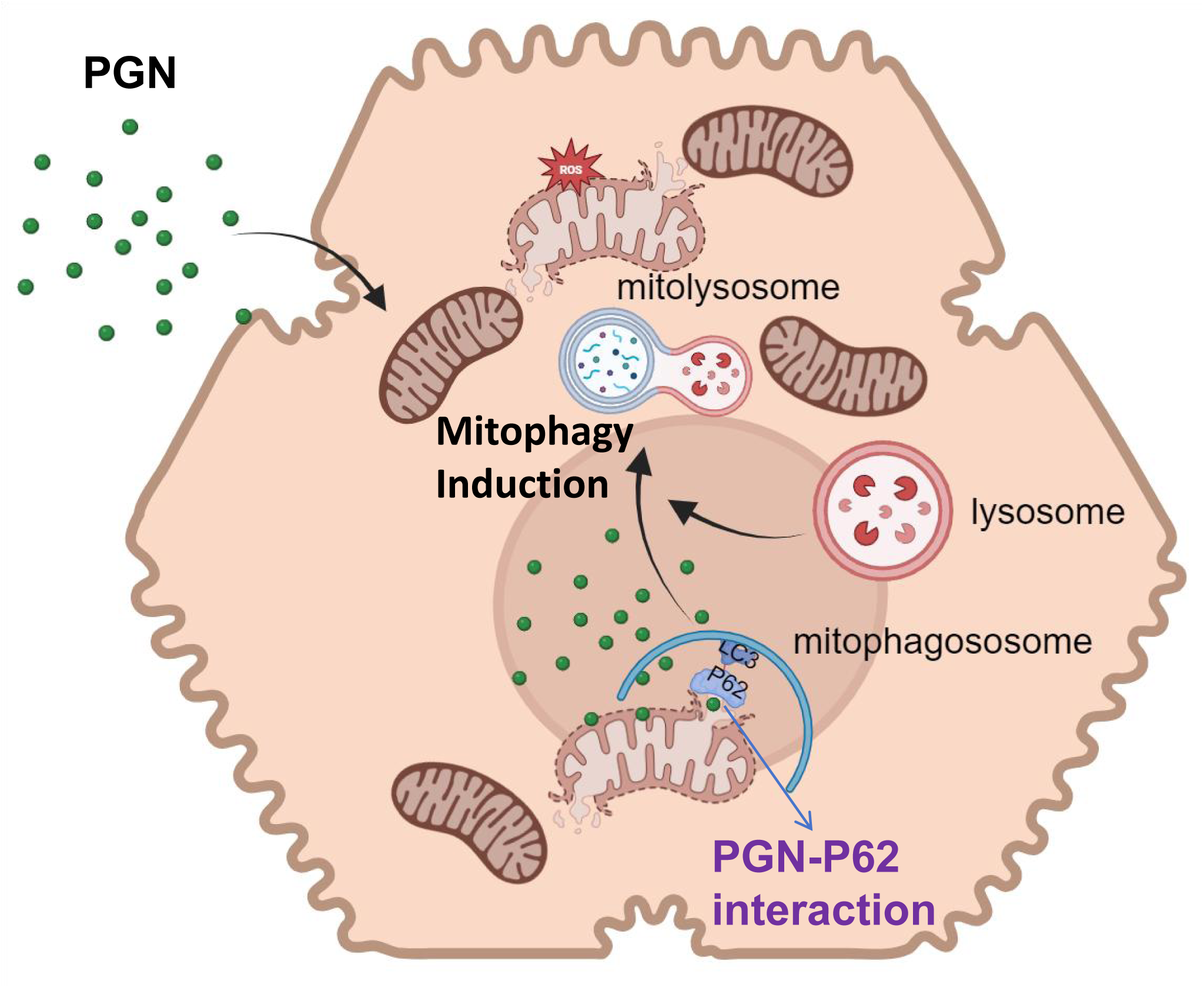
PGN-p62 Axis Activates Mitophagy to Protect Against Hepatocyte Death. Bacterial PGN activates hepatocyte mitophagy via p62 binding, improving mitochondrial homeostasis and reducing hepatocyte death under stress.

Beyond its well-established role in host immunity(Wolf & Underhill, 2018), our study reveals that PGN functions as a critical regulator of mitophagy—a finding that expands the understanding of microbial influences on cellular homeostasis. Intriguingly, the ability of PGN to modulate mitochondrial health aligns with the endosymbiotic theory of mitochondrial evolution, which posits that mitochondria originated from ancient bacteria (Roger *et al*, 2017). Given that mitochondria share structural and functional similarities with bacteria (Gogoi *et al*, 2022), it is plausible that bacterial cell wall components, such as PGN, may have co-evolved to interact with mitochondrial quality control mechanisms. This discovery suggests that host-microbe coevolution has fine-tuned PGN as a conserved signal for mitochondrial surveillance, highlighting a fundamental link between bacterial-derived molecules and organelle homeostasis.

Although the gut harbors vast quantities of bacteria (de Vos *et al*, 2022), our findings suggest that exogenous PGN may still be required to mitigate liver injury. One possible explanation is that liver disease disrupts gut microbiota composition, particularly reducing Gram-positive bacteria that are a major source of PGN (Pasquina-Lemonche *et al*, 2020), thereby creating a deficit in PGN availability. Alternatively, under conditions of liver damage, the host may prioritize energy allocation toward hepatic repair, leading to reduced secretion of gut lysozymes that normally degrade bacterial cell walls and release PGN (Ragland & Criss, 2017). Consequently, systemic PGN levels may decline precisely when hepatoprotective mitophagy is most needed. These hypotheses warrant further investigation into how liver-gut crosstalk modulates PGN availability during disease.

The progression from hepatocyte death to hepatic fibrosis follows a well-characterized pathological sequence: damaged hepatocytes release damage-associated molecular patterns (DAMPs) and pro-inflammatory mediators, which in turn activate HSCs, leading to excessive extracellular matrix deposition and fibrotic scarring (Luedde *et al*, 2014). Our study demonstrates that PGN alleviates hepatocyte death by inducing p62-dependent mitophagy. By selectively clearing dysfunctional mitochondria—the primary source of ROS and pro-apoptotic factors in CCl4-induced injury (Liu *et al*, 2018) - PGN reduces mitochondrial stress and attenuates hepatocyte necrosis. This protective effect subsequently diminishes the activation of HSCs and collagen deposition, thereby interrupting the progression from hepatocyte injury to fibrosis. Notably, PGN’s action mechanism exhibits evolutionary refinement, as it promotes mitochondrial quality control without inducing excessive autophagy - a critical advantage over pan-autophagy activators that may provoke uncontrolled cellular catabolism (Levine *et al*, 2015).

These findings position PGN-mediated mitophagy as a promising therapeutic strategy not only for liver fibrosis but potentially for other diseases characterized by mitochondrial dysfunction. The evolutionary conservation of PGN-mitochondria interactions suggests this mechanism may represent a fundamental host-microbe adaptation for organelle homeostasis. While our data demonstrate efficacy in chronic liver injury models, future studies should investigate whether PGN maintains its protective effects in other conditions featuring sustained mitochondrial damage, such as non-alcoholic steatohepatitis (Tian *et al*, 2022) or drug-induced liver injury (Wang *et al*, 2024). Moreover, given the central role of mitochondrial quality control in neurodegenerative diseases (Cheng *et al*, 2022) and metabolic syndromes (Ng *et al*, 2021), exploring PGN’s therapeutic potential in these contexts could yield significant clinical benefits. This approach would capitalize on an ancient symbiotic relationship to address modern diseases of mitochondrial dysfunction.

In summary, our work identifies bacterial PGN as a novel guardian of mitochondrial integrity, acting through a p62-dependent mitophagy pathway to protect against liver injury. These findings deepen our understanding of how microbial molecules influence host cell physiology and open new avenues for developing microbiome-inspired therapies targeting mitochondrial quality control.

## Resource availability

### Lead contact

Further information and requests for reagents may be directed to the Lead contact Zhao Shan.

### Materials availability

All reagents and strains generated by this study are available through request to the lead contact with a completed Material Transfer Agreement.

### Data and code availability

RNA-seq data are provided in Table S2

This paper does not report original code.

Any additional information required to reanalyze the data reported in this paper is available from the lead contact upon request.

## Acknowledgements

We thank Dr. Yonglong Wei (Yunnan University) for providing Cas9 mice; Dr. Yuanxiang Zhu (Yunnan University) for helping in imaging analysis; Dr. Xuna Wu (Yunnan University) for helping in Mass Spectrometry; Dr. Hao Yin (Wuhan University) for providing AML12 cells; Dr. Qiang Ding (Qinghua University) for providing Huh7 cells; Dr. Mei Yang (Yunnan University) for providing HEK293T cells. This work was supported by Yunnan Provincial Science and Technology Project at Southwest United Graduate School (202302AP370005 to B.Q.), National Natural Science Foundation of China (32170794 to B.Q.; 32071129 to Z.S.), Yunnan Revitalization Talent Support Program (C619300A086 to Z.S., K264202230211 to B.Q.).

## Author Contributions

J.T., L.W., X.W. performed experiments, analyzed data, and J.T. wrote the figure legends. M.T. wrote the methods. L.W. revised the figure legends and methods. B.Q and Z.S. conceived this study, analyzed data, and wrote the paper.

## STAR★Methods

### Experimental model and study participant details

#### Mice

C57BL/6J mice (strain #N000013) were obtained from GemPharmatech. Rosa26-Cas9 knockin mice (The Jackson Laboratory strain #024858) were provided by Dr. Yonglong Wei (Yunnan University). All mice were housed under standard conditions in the animal core facility of Yunnan University. Animal protocols were approved by the Institutional Animal Care and Use Committee (IACUC) of Yunnan University (No. YNU20220262).

### Method details

#### Acetaminophen (APAP)-Induced Liver Injury (AILI)

Male mice (8–10 weeks old) were fasted overnight (5:00 p.m. to 9:00 a.m.) before intraperitoneal (i.p.) injection of 210 mg/kg APAP (Sigma, A7085). Immediately after APAP administration, mice received oral gavage of either heat-killed E. coli OP50 (HK OP50, 16 mg/mouse in 200 µL ddH₂O) or ddH₂O (control). Liver tissues and sera were collected at indicated time points. Liver injury was assessed by H&E staining and serum alanine aminotransferase (ALT) levels using a diagnostic assay kit (Teco Diagnostics).

#### CCl₄-Induced Liver Injury

Male mice (8–10 weeks old) were injected i.p. with 10% CCl₄ (in corn oil) twice weekly (initial dose: 4 µL/g body weight; subsequent doses: 2 µL/g). Control mice received corn oil. From week 3 onward, mice were orally gavaged with E. coli OP50 peptidoglycan (PGN, 16 mg/mouse, 3×/week) or ddH₂O (control). Liver tissues and sera were harvested as described.

#### Mitochondrial Keima (mtKeima) Imaging

Male mice (6 weeks old, 18–20 g) were injected via tail vein with AAV8-TBG-mtKeima (1×10¹¹ vg/mouse). For P62 knockout studies, Rosa26-Cas9 mice were co-injected with AAV8-TBG-mtKeima and AAV8-TBG-sgCtrl/sgP62 (1×10¹² vg/mouse). Two weeks post-injection, liver tissues were sectioned (1 mm) and imaged within 30 min post-euthanasia using a Zeiss LSM 900 confocal microscope (561 nm/405 nm dual-excitation, 647 nm emission).

### Cell Culture and Treatments

#### Cell Lines, Primary Hepatocytes Isolation and Maintenance

HEK293T, Huh7, and AML12 cells were cultured in DMEM (VivaCell, #C3113-0500) supplemented with 10% FBS (VivaCell, #C04001-500), 100 IU/mL penicillin, and 100 μg/mL streptomycin (VivaCell, #C3420-0100) at 37°C under 5% CO₂. All cell lines were routinely confirmed mycoplasma-free by qRT-PCR. Primary hepatocytes were isolated as previously described(Shan *et al*, 2018).

#### Transfection

Cells were transfected using polyethyleneimine (PEI; Biohub, #49553-93-7) at a 1:3 plasmid-to-PEI ratio (w/v), with total DNA normalized using empty vector controls. Cells and supernatants were harvested 48 hours post-transfection for downstream assays.

#### Pharmacological and PGN Treatments

##### Mdivi-1 Treatment

Cells at 50% confluency were pretreated with 20 μM or 50 μM Mdivi-1 (Selleck, #HY-101461) or DMSO vehicle control for 4 hours, followed by stimulation with 1mg *E. coli* PGN or PBS (100 µL) for 24 hours. A second dose of Mdivi-1 or DMSO was administered 4 hours before harvest.

##### APAP/PGN Treatment

Huh7 or AML12 cells seeded on glass coverslips (70% confluency) were treated with 2 mM (Huh7) or 5 mM (AML12) APAP plus PBS or PGN (1mg) for 24–48 hours.

#### Live-Cell Imaging

##### Mitochondrial Staining

AML12 cells were stained with 100 nM MitoTracker Green (Thermo, #M7514) and 200 nM MitoTracker Deep Red (Thermo, #M22426) for 15 minutes at 37°C and imaged using a Zeiss LSM 900 confocal microscope (63× oil objective).

##### SYTOX Green Assay

Cells were incubated with 400 nM SYTOX Green (Thermo, #S7020) for 20 minutes post-PGN treatment, followed by PBS washes.

#### Immunofluorescence on cells

Cells were fixed with 4% PFA (15 min, RT), permeabilized with 0.2% Triton X-100 (10 min, RT), and blocked with 5% goat serum + 2.5% BSA (1 hr, RT). Primary antibodies against Ki67 (Abcam, #ab15580, 1:200), TOMM20 (Thermo, #11802-1-AP, 1:200), LAMP1 (DSHB, #AB_528127, 1:200), P62 (ABclonal, #A19700, 1:200), or LC3 (MBL, #PM036, 1:200) were incubated overnight at 4°C. Alexa Fluor 488/568-conjugated secondary antibodies (Jackson ImmunoResearch/Thermo, 1:800) and DAPI (Beyotime, #C1006) were used for detection. Images were acquired using a Zeiss LSM 900 confocal microscope.

#### Lentiviral Production and Stable Cell Line Generation

Lentivirus was produced in HEK293T cells (≥70% confluency) by co-transfecting the transfer plasmid (pHage-TO-mtKeima) with packaging plasmids psPAX2 and pVSVG (4:3:1 ratio) using PEI/Opti-MEM complexes. After 15 min incubation, the mixture was added to cells, and medium was replaced within 12–18 h. Viral supernatants were collected at 48 h, filtered (0.22 µm), and used to infect AML12 or Huh7 cells (1 mL/well). Stable transductants were selected with puromycin (1 µg/mL for AML12, 5 µg/mL for Huh7) for 7 days.

#### AAV Production and Purification

AAV vectors were packaged in HEK293T cells (passages P2-P10) cultured in 175 cm² flasks at >70% confluency. For transfection, plasmids (target vector and RC8 helper plasmid) were mixed with Opti-MEM (3 mL) at a concentration calculated by DNA size (1 bp = 0.000983 µg), followed by addition of PEI at a 1:3 plasmid:PEI ratio. After 15 min incubation, the mixture was added to cells. At 24 h post-transfection, medium was replaced with serum-free DMEM containing 1% penicillin/streptomycin. Cells and supernatants were harvested 72 h later for purification.

Viral particles were purified through sequential steps: (1) Chloroform extraction (3 mL) and NaCl precipitation (7.6 mL of 5M) followed by centrifugation (4,000 ×g, 8 min, 4°C); (2) PEG8000 precipitation (9.4 mL of 50% solution, 1 h on ice) and centrifugation (4,000 ×g, 30 min); (3) Pellet resuspension in 50 mM HEPES (pH 8.0) with MgCl₂ (2.5 mM), DNase I (10 U/mL), and RNase A (10 µg/mL) at 37°C for 20 min; (4) Double chloroform extraction (1:1 ratio) and centrifugation (3,000 ×g, 5 min); (5) Final concentration using ultrafiltration (14,000 ×g washes with PBS until solution cleared).

Viral titers were determined by qPCR. Viral DNA was digested with DNase/proteinase K, and quantified against a plasmid standard curve (1-0.0078 ng/µL). Reactions contained SYBR Green Master Mix, primers (5 µM), and template DNA in 15 µL total volume.

#### Flow Cytometric Analysis of Mitophagy and Mitochondrial Function Mitophagy Assessment (mtKeima)

Huh7 cells stably expressing mtKeima were treated with PGN or DMEM control (containing 2 µg/mL puromycin) for 24 hours. Following trypsinization and PBS washes, cells were resuspended in PBS with 2% FBS. CCCP (50 µM, 2 hours; Sigma #C2759) served as a positive control. mtKeima fluorescence was analyzed using a BD FACSAria III flow cytometer with dual-excitation (405 nm for neutral pH, 561 nm for acidic pH) and emission detection (614-620 nm). A minimum of 10,000 events per sample were recorded, with data analysis performed using FlowJo 10.4.

#### Mitochondrial Membrane Potential (ΔΨm) Measurement

HEK293T cells were treated with PGN or PBS for 24 hours, harvested by trypsinization, and stained with 200 nM TMRM for 30 minutes at 37°C/5% CO₂. Following PBS washes, ΔΨm was assessed on a BD FACSAria III (488 nm excitation/570 nm emission). APAP (5 mM, 24 hours) served as a positive control for mitochondrial depolarization. Data were gated based on FSC/SSC parameters and TMRM fluorescence (n=3 biological replicates per group).

#### Histology and Immunofluorescence Staining

##### Hematoxylin and Eosin (H&E) and Sirius Red Staining

Liver tissue paraffin sections were stained with H&E. Briefly, liver tissue was fixed overnight in 10% formalin, dehydrated through a graded alcohol series, and embedded in paraffin before 5 µm sectioning. H&E staining was performed using Hematoxylin (Biocare Medical, cat# A600701-0050) and Eosin (Edgar Degas Eosin, Biocare Medical, THE-MM, cat# 55621322). Liver tissue paraffin sections were stained using a Sirius Red staining kit (Solarbio, G1472) according to the manufacturer’s protocol to evaluate collagen deposition and fibrosis extent.

##### Immunofluorescence Staining on Frozen Sections

Fresh liver tissue was immediately fixed in 4% paraformaldehyde for 1 hour. After three washes in PBS, the tissue was dehydrated overnight in 30% sucrose at 4°C and then embedded in OCT. The tissue was cut into 5 μm thick frozen sections and fixed in pre-cooled acetone at −20°C for 15 minutes. The sections were permeabilized with 0.2% Triton X-100 in PBS and blocked with blocking solution (5% goat serum + 0.3% BSA in PBS). The sections were then incubated overnight at 4°C with primary antibody, anti-Desmin antibody (Proteintech, Cat #16520-1-AP, 1:500). After washing with TBST, the sections were incubated for 1 hour at room temperature with Alexa Fluor 488-conjugated Affinipure Goat Anti-Rabbit secondary antibody (Jackson ImmunoResearch, Cat # 111-545-003, 1:800). After TBST washing, the sections were stained with DAPI (Beyotime, Cat #C1006) for nuclear staining, mounted, and imaged using a Zeiss LSM 900 confocal microscope. All images were captured under consistent parameters.

#### Flow Cytometric Analysis of Mitophagy and Mitochondrial Membrane Potential (ΔΨm)

##### Mitophagy Assay (mtKeima)

Huh7 cells stably expressing mtKeima were treated with PGN or DMEM control in 2 µg/mL puromycin for 24 hours. After trypsinization, cells were washed with PBS and resuspended in PBS containing 2% FBS. CCCP (Sigma #C2759, [50 µM, 2h]) was used as a positive control. MtKeima fluorescence was measured on a BD FACSAria III with dual excitation (405 nm for neutral pH, 561 nm for acidic pH) and emission detection (620/29 nm, 614/20 nm). At least 10,000 events/sample were recorded, and data were analyzed using FlowJo 10.4.

##### Mitochondrial Membrane Potential (ΔΨm, TMRM Assay)

HEK293T cells were treated with PBS or PGN for 24 hours, harvested by trypsinization, and neutralized with DMEM/10% FBS. After centrifugation (1000 rpm, 3 min), cells were stained with 200 nM TMRM for 30 min at 37°C/5% CO₂, washed twice with PBS, and resuspended in PBS/2% FBS. APAP ([5mM, 24h]) served as a positive control for mitochondrial depolarization. ΔΨm was assessed on a BD FACSAria III using the PE channel (488 nm excitation/570 nm emission). Data were gated based on FSC/SSC vs. TMRM fluorescence, with n=3/group replicates.

#### Histology and Immunofluorescence Staining

##### Hematoxylin and Eosin (H&E) and Sirius Red Staining

Liver tissue paraffin sections were stained with H&E. Briefly, liver tissue was fixed overnight in 10% formalin, dehydrated through a graded alcohol series, and embedded in paraffin before 5 µm sectioning. H&E staining was performed using Hematoxylin (Biocare Medical, cat# A600701-0050) and Eosin (Edgar Degas Eosin, Biocare Medical, THE-MM, cat# 55621322). Liver tissue paraffin sections were stained using a Sirius Red staining kit (Solarbio, G1472) according to the manufacturer’s protocol to evaluate collagen deposition and fibrosis extent.

##### Immunofluorescence Staining on frozen liver sections

Fresh liver tissue was immediately fixed in 4% paraformaldehyde for 1 hour. After three washes in PBS, the tissue was dehydrated overnight in 30% sucrose at 4°C and then embedded in OCT. The tissue was cut into 5 μm thick frozen sections and fixed in pre-cooled acetone at −20°C for 15 minutes. The sections were permeabilized with 0.2% Triton X-100 in PBS and blocked with blocking solution (5% goat serum + 0.3% BSA in PBS). The sections were then incubated overnight at 4°C with primary antibody, anti-Desmin antibody (BioLegend, Proteintech, Cat #16520-1-AP, 1:500). After washing with TBST, the sections were incubated for 1 hour at room temperature with Alexa Fluor 647-conjugated Affinipure Goat Anti-Rabbit secondary antibody (Jackson ImmunoResearch, Cat # 115-605-003, 1:800). After TBST washing, the sections were stained with DAPI (Beyotime, Cat #C1006) for nuclear staining, mounted, and imaged using a Zeiss LSM 900 confocal microscope. All images were captured under consistent parameters.

#### Preparation of Heat-Inactivated *E. coli* and PGN

A single colony of *E. coli* OP50 (CGC, WBStrain00041969) was cultured overnight in 10 mL LB broth (10 g/L tryptone [Sigma,cat#LP0042], 5 g/L yeast extract [Sigma,cat# LP0021B], 5 g/L NaCl [GHTECH,cat# 1266-2006]) with shaking until reaching an OD600 of ∼1.2. Bacterial suspensions were heat-inactivated (85°C, 45 min), pelleted (8,000 ×g, 5 min), and resuspended in PBS. Complete inactivation was confirmed by absence of colony growth on LB agar plates.

PGN was isolated as previously described (Hao *et al*., 2024). Briefly, overnight cultures of *E. coli* OP50 (60 mL LB) were pelleted (8000 ×g, 5 minutes) and resuspended in 6 mL 1M NaCl solution ([1.168g NaCl GHTECH, cat#1266-2006, 20ml ddH₂O]). The suspension was boiled (100°C, 30 minutes), washed 3× with ddH₂O, and sonicated on ice for 1 hour. The lysate was centrifuged (10,000 ×g, 5 minutes), and the pellet was digested sequentially in: Solution B (5mg/ml 150µl DNase, 5mg/ml 180µl RNase,14.67ml PH 6.8 Tris-HCL cat#T1503) at 37°C for 1 hour; 50 µg/mL trypsin (Sangon, cat#A100260-0050) at 37°C for 1 hour. After 3× ddH₂O washes, the sample was reboiled (100°C, 5 minutes). Purified PGN was prepared by centrifugation (8,000 × g, 5 min) and quantified gravimetrically prior to use. PGN was labeled with FITC as previously described (Mann *et al*, 2016). Briefly, 500mg PGN was labeled with 1 mg/mL FITC (Sigma cat# F7250) in the dark (1 hour, RT). Excess FITC was removed by 3× PBS washes. Labeling efficiency was assessed by microscopy.

#### Molecular Cloning & Protein Interaction Analysis

The coding sequences of LC3 and P62 were cloned into pIPMyc/Flag or pmCherry vectors using restriction enzyme-based methods. For exogenous interaction studies, we transfected HEK293T cells with various tagged constructs (mCherry-, Myc-, or Flag-tagged P62 variants or LC3). Following lysis in RIPA buffer, cell extracts were incubated with 100 μg peptidoglycan (PGN) overnight at 4°C with rotation. The PGN-protein complexes were subsequently pelleted by centrifugation (12,000 × g, 10 min, 4°C) and washed three times with PBS. For endogenous interaction studies, AML12 cells were treated with either PBS (control) or 100 μg PGN for 48 hours prior to lysis. Endogenous P62-PGN complexes were isolated by centrifugation, while P62-associated proteins were immunoprecipitated using anti-P62 antibody (1 μg; ABclonal #A19700) or species-matched IgG control bound to Protein A/G magnetic beads (6 μL/sample; Thermo #88802) for 2 hours at 4°C. All precipitates were washed five times with 500 μL PBS and bound proteins were eluted in 50 μL PBS, denatured in SDS loading buffer, and analyzed by SDS-PAGE and Western blotting.

#### Prokaryotic Expression and Interaction Validation

The P62 coding sequence was cloned into the pET28a vector and expressed in E. coli BL21(DE3) cells. Protein expression was induced with 1 mM IPTG at 18°C overnight, yielding high levels of His-tagged P62. The recombinant protein was purified under native conditions using Ni-NTA affinity chromatography according to the manufacturer’s protocol. Bound proteins were eluted with buffer containing 500 mM imidazole (pH 8.0), and purity was confirmed by SDS-PAGE and Coomassie Brilliant Blue staining. Protein concentration was determined using a BCA assay.

For microscopy-based interaction studies, His-P62-bound Ni-NTA beads (2 μg protein + 50 μL beads) were incubated with FITC-labeled PGN (200 μg) overnight at 4°C. Following two washes with HEPES buffer (25 mM HEPES pH 7.5, 150 mM NaCl, 1 mM DTT), samples were imaged using an upright fluorescence microscope (Leica DM6 B) with brightfield and GFP filter sets. Control experiments included Elution Buffer-bound beads and unlabeled PGN samples.

#### Western Blot Analysis

Cells were lysed on ice for 30 minutes using RIPA buffer (10 mM Tris-HCl pH 8.0, 1 mM EDTA, 0.5 mM EGTA, 1% Triton X-100, 0.1% sodium deoxycholate, 0.1% SDS, 140 mM NaCl) freshly supplemented with 100 mM PMSF and protease inhibitors (BBI, cat# A610425-0025). Lysates were centrifuged (12,000 × g, 10 min, 4°C), and supernatants were resolved by SDS-PAGE (12.5% gel) and transferred to PVDF/nitrocellulose membranes (0.45 μm). Membranes were blocked with 5% 5% non-fat milk for 1 hour at RT, then incubated overnight at 4°C with primary antibodies: Cyp2e1 (LSBio, #LS-C6332, 1:1000); NAPQI (Provided by Cynthia Ju Lab, UTHealth, 1:800); SQSTM1/p62 (ABclonal, #A19700, 1:20,000); LC3 (MBL, #PM036, 1:1000); β-Actin (Proteintech, #66009-1, 1:20,000). After HRP-conjugated secondary antibody (Thermo, Goat anti-mouse IgG (H+L) Secondary antibody, #626520,1:8000; Invitrogen, Goat anti-Rabbit IgG (H+L) Secondary Antibody,# 65-6140) incubation, proteins were detected using ECL Select (Cytiva, #RPN2235) or SuperSignal West Pico PLUS (Thermo, #34580).

#### Transmission Electron Microscopy (TEM)

AML12 cells grown on coverslips in 6-well plates were fixed sequentially under the following conditions: Primary fixation: 2.5% glutaraldehyde in culture medium (5 min, RT), followed by fixation in 2.5% glutaraldehyde in 0.1 M PBS (4°C, 2 h). Post-fixation: 1% osmium tetroxide (OsO₄) in 0.1 M PBS (1 h, 4°C). After fixation, samples were dehydrated through a graded ethanol series (30%, 50%, 70%, 90%, and 100%, 10 min per step) and infiltrated with [35ml epoxy resin (15ml SPI Supplies, SPI-812#02659-AB, 10ml DDSA#02827-AF,10ml NMA#02828-AF,660µl DMP-30#02823-DA] in incremental concentrations (25%, 50%, 75%, and 100%, 2 h per step). The resin was polymerized at 60°C for 48 h. Ultrathin sections (70 nm) were cut using an ultramicrotome, stained with uranyl acetate (10 min) and lead citrate (5 min), and imaged on a Leica EM UC7 transmission electron microscope. For quantification of mitochondrial morphology and number via transmission electron microscopy (TEM), TEM images were first imported into ImageJ software (v1.54p). Scale calibration was performed using the nanometer-scale ruler embedded in the images to convert pixel units to physical units. The boundary of the target cell was then manually delineated to exclude extracellular matrix and adjacent cells. Mitochondria were identified based on their characteristic ultrastructural features, including double membranes, internal cristae, and moderately electron-dense matrices, with strict differentiation from organelles such as lysosomes and peroxisomes during counting. For each cell, mitochondrial particles were manually labeled in multiple regions (e.g., 5 randomly selected representative fields of view), and the total number of mitochondria within the entire cell was aggregated.

#### RNA Sequencing

Total RNA was extracted using RNAiso Plus (TaKaRa, #9109) and assessed for purity (NanoDrop 2000) and integrity (Agilent Bioanalyzer 2100, RNA Nano 6000 Kit). Libraries were prepared from 1 μg RNA using the NEB Next Ultra RNA Library Prep Kit and sequenced on an Illumina NovaSeq 6000 (paired-end, 150 bp, BioMarker). Raw reads were filtered (The Q30 base percentage ranged from 93.39% (PBS1) to 94.21% (APPGN1), and the GC content varied from 48.91% (APPGN2) to 49.83% (PBS2). and aligned to GRCm38 (Hisat2). Differential expression (limma) and enrichment (clusterProfiler) analyses were performed in RStudio (v4.3).

#### Liver Tissue Processing and LC-MS/MS Analysis

Liver tissue samples (30 mg wet weight) were homogenized in ice-cold lysis buffer (0.1 M Tris-HCl, 1% sodium deoxycholate, pH 7.4) supplemented with complete protease inhibitor cocktail (MCE, Cat# HY-K0011). The homogenate was centrifuged at 10,000 ×g for 10 min at 4°C, and the resulting supernatant was incubated with 300 mg PGN overnight at 4°C with gentle rotation. PGN-protein complexes were pelleted by centrifugation (12,000 ×g, 10 min, 4°C) and washed four times with 50 mM MES buffer (pH 6.0, containing 150 mM NaCl). PGN-binding proteins were identified by LC-MS/MS using a Q Exactive HF-X mass spectrometer (Thermo Scientific) at Yunnan University’s Core Facility. Glutathione (GSH) quantification was performed by LC-MS/MS (SCIEX TripleTOF 6600 system) using authentic GSH standard (Sigma-Aldrich, Cat# G4251) for calibration.

#### Data and Statistical Analysis

All experimental data are presented as mean ± standard error of the mean (SEM). Statistical analyses were performed using GraphPad Prism (v8.0.2), with appropriate tests selected based on experimental design: unpaired two-tailed Student’s t-tests for comparisons between two groups, and one-way ANOVA for comparisons involving three or more groups. For image-based quantifications, regions of interest (ROIs) were systematically selected and analyzed using ImageJ software. Fluorescence intensity profiles were generated by measuring pixel gray values along predefined lines within magnified ROIs, with distinct color-coded curves (red/green) representing different fluorescence channels (e.g., MTDR/Mtkeima PH4 vs MTG/Mtkeima PH7). Multiple ROIs and measurement lines were analyzed per condition to ensure comprehensive characterization of fluorescence distribution patterns.

Colocalization studies were conducted by selecting ROIs containing target cells from merged fluorescence images (e.g., Tomm20/P62 or P62/LC3 dual-labeled samples). The Coloc2 plugin in ImageJ was employed to extract paired channel pixel intensities and generate two-dimensional intensity histograms. Colocalization strength was quantified using Pearson’s correlation coefficient (R), with a minimum of five cells analyzed per condition to establish statistical validity. This rigorous analytical approach ensured robust quantification of protein interactions and subcellular localization patterns across all experimental conditions.

## References

Bock FJ, Tait SWG (2020) Mitochondria as multifaceted regulators of cell death. Nat Rev Mol Cell Bio 21: 85–100

Chen JX, Jian LE, Guo YK, Tang CW, Huang ZY, Gao JH (2024) Liver Cell Mitophagy in Metabolic Dysfunction-Associated Steatotic Liver Disease and Liver Fibrosis. Antioxidants-Basel 13

Cheng XT, Huang N, Sheng ZH (2022) Programming axonal mitochondrial maintenance and bioenergetics in neurodegeneration and regeneration. Neuron 110: 1899–1923

Chimerel C, Emery E, Summers DK, Keyser U, Gribble FM, Reimann F (2014) Bacterial Metabolite Indole Modulates Incretin Secretion from Intestinal Enteroendocrine L Cells. Cell Reports 9: 1202–1208

Chimerel C, Murray AJ, Oldewurtel ER, Summers DK, Keyser UF (2013) The Effect of Bacterial Signal Indole on the Electrical Properties of Lipid Membranes. Chemphyschem 14: 417–423

de Vos WM, Tilg H, Van Hul M, Cani PD (2022) Gut microbiome and health: mechanistic insights. Gut 71: 1020–1032

Engelmann C, Clària J, Szabo G, Bosch J, Bernardi M (2021) Pathophysiology of decompensated cirrhosis: Portal hypertension, circulatory dysfunction, inflammation, metabolism and mitochondrial dysfunction. Journal of Hepatology 75: S49–S66

Gogoi J, Bhatnagar A, Ann KJ, Pottabathini S, Singh R, Mazeed M, Kuncha SK, Kruparani SP, Sankaranarayanan R (2022) Switching a conflicted bacterial DTD-tRNA code is essential for the emergence of mitochondria. Science Advances 8

Hao FR, Liu HM, Qi B (2024) Bacterial peptidoglycan acts as a digestive signal mediating host adaptation to diverse food resources in C. elegans. Nature Communications 15

Hsu CL, Schnabl B (2023) The gut-liver axis and gut microbiota in health and liver disease. Nat Rev Microbiol 21: 719–733

Hu MY, Xu Y, Wang YQ, Huang ZH, Wang L, Zeng FN, Qiu BW, Liu ZF, Yuan PB, Wan Y et al (2025) Gut microbial-derived N-acetylmuramic acid alleviates colorectal cancer via the AKT1 pathway. Gut

Le HH, Lee MT, Besler KR, Johnson EL (2022) Host hepatic metabolism is modulated by gut microbiota-derived sphingolipids. Cell Host Microbe 30: 798-+

Levine B, Packer M, Codogno P (2015) Development of autophagy inducers in clinical medicine. Journal of Clinical Investigation 125: 14–24

Lin ZF, Wu F, Lin SQ, Pan XB, Jin LG, Lu TT, Shi LH, Wang Y, Xu AM, Li XK (2014) Adiponectin protects against acetaminophen-induced mitochondrial dysfunction and acute liver injury by promoting autophagy in mice. Journal of Hepatology 61: 825–831

Liu PF, Liu XY, Qi B (2024) UPR-immunity axis acts as physiological food evaluation system that promotes aversion behavior in sensing low-quality food. Elife 13

Liu Y, Wen PH, Zhang XX, Dai Y, He Q (2018) Breviscapine ameliorates CCl-induced liver injury in mice through inhibiting inflammatory apoptotic response and ROS generation. Int J Mol Med 42: 755–768

Luedde T, Kaplowitz N, Schwabe RF (2014) Cell Death and Cell Death Responses in Liver Disease: Mechanisms and Clinical Relevance. Gastroenterology 147: 765–U110

Ma XW, Niu MW, Ni HM, Ding WX (2024) Mitochondrial dynamics, quality control, and mtDNA in alcohol-associated liver disease and liver cancer. Hepatology

Mann B, Loh LN, Gao G, Tuomanen E (2016) Preparation of Purified Gram-positive Bacterial Cell Wall and Detection in Placenta and Fetal Tissues. Bio-protocol 6

Mann ER, Lam YK, Uhlig HH (2024) Short-chain fatty acids: linking diet, the microbiome and immunity. Nature Reviews Immunology 24: 577–595

Mansouri A, Gattolliat CH, Asselah T (2018) Mitochondrial Dysfunction and Signaling in Chronic Liver Diseases. Gastroenterology 155: 629–647

Minton K (2015) Mitochondria and phagosomes: better together. Nature Reviews Immunology 15: 667–667

Ng MYW, Wai T, Simonsen A (2021) Quality control of the mitochondrion. Developmental Cell 56: 881–905

Palikaras K, Lionaki E, Tavernarakis N (2018) Mechanisms of mitophagy in cellular homeostasis, physiology and pathology. Nat Cell Biol 20: 1013–1022

Pasquina-Lemonche L, Burns J, Turner RD, Kumar S, Tank R, Mullin N, Wilson JS, Chakrabarti B, Bullough PA, Foster SJ et al (2020) The architecture of the Gram-positive bacterial cell wall. Nature 582: 294-+

Ragland SA, Criss AK (2017) From bacterial killing to immune modulation: Recent insights into the functions of lysozyme. Plos Pathogens 13

Roger AJ, Muñoz-Gómez SA, Kamikawa R (2017) The Origin and Diversification of Mitochondria. Current Biology 27: R1177–R1192

Schwabe RF, Luedde T (2018) Apoptosis and necroptosis in the liver: a matter of life and death. Nat Rev Gastro Hepat 15: 738–752

Shan Z, Liu X, Chen Y, Wang M, Gao YR, Xu L, Dar WA, Lee CG, Elias JA, Castillo PD et al (2018) Chitinase 3-like-1 promotes intrahepatic activation of coagulation through induction of tissue factor in mice. Hepatology 67: 2384–2396

Sun YQ, Cao Y, Wan HY, Memetimin A, Cao Y, Li L, Wu CY, Wang M, Chen S, Li Q et al (2024) A mitophagy sensor PPTC7 controls BNIP3 and NIX degradation to regulate mitochondrial mass. Mol Cell 84

Tian C, Min XW, Zhao YX, Wang YC, Wu XS, Liu ST, Dou W, Zhou TT, Liu Y, Luo RK et al (2022) MRG15 aggravates non-alcoholic steatohepatitis progression by regulating the mitochondrial proteolytic degradation of TUFM. Journal of Hepatology 77: 1491–1503

Tian D, Cui MX, Han M (2024) Bacterial muropeptides promote OXPHOS and suppress mitochondrial stress in mammals. Cell Reports 43

Tian D, Han M (2022) Bacterial peptidoglycan muropeptides benefit mitochondrial homeostasis and animal physiology by acting as ATP synthase agonists. Developmental Cell 57: 361-+

Tujios S, Fontana RJ (2011) Mechanisms of drug-induced liver injury: from bedside to bench. Nat Rev Gastroenterol Hepatol 8: 202–211

Wang JJ, Qiu YP, Yang LJ, Wang JC, He J, Tang CW, Yang ZX, Hong WX, Yang B, He QJ et al (2024) Preserving mitochondrial homeostasis protects against drug-induced liver injury via inducing OPTN (optineurin)-dependent Mitophagy. Autophagy 20: 2677–2696

Wolf AJ, Underhill DM (2018) Peptidoglycan recognition by the innate immune system. Nature Reviews Immunology 18: 243–254

Yang WJ, Cong YZ (2021) Gut microbiota-derived metabolites in the regulation of host immune responses and immune-related inflammatory diseases. Cellular & Molecular Immunology 18: 866–877

Yin RP, Wang T, Sun JZ, Dai HQ, Zhang YT, Liu NN, Liu HW (2025) Postbiotics From Lactobacillus Johnsonii Activates Gut Innate Immunity to Mitigate Alcohol-Associated Liver Disease. Adv Sci 12

Zhang IW, Curto A, López-Vicario C, Casulleras M, Duran-Güell M, Flores-Costa R, Colsch B, Aguilar F, Aransay AM, Lozano JJ et al (2022) Mitochondrial dysfunction governs immunometabolism in leukocytes of patients with acute-on-chronic liver failure. Journal of Hepatology 76: 93–106

