## Supplemental Figures for "Microbial Peptidoglycan Engages Autophagy Receptor P62 to Induce Protective Mitophagy in the Liver"

**Figure S1**

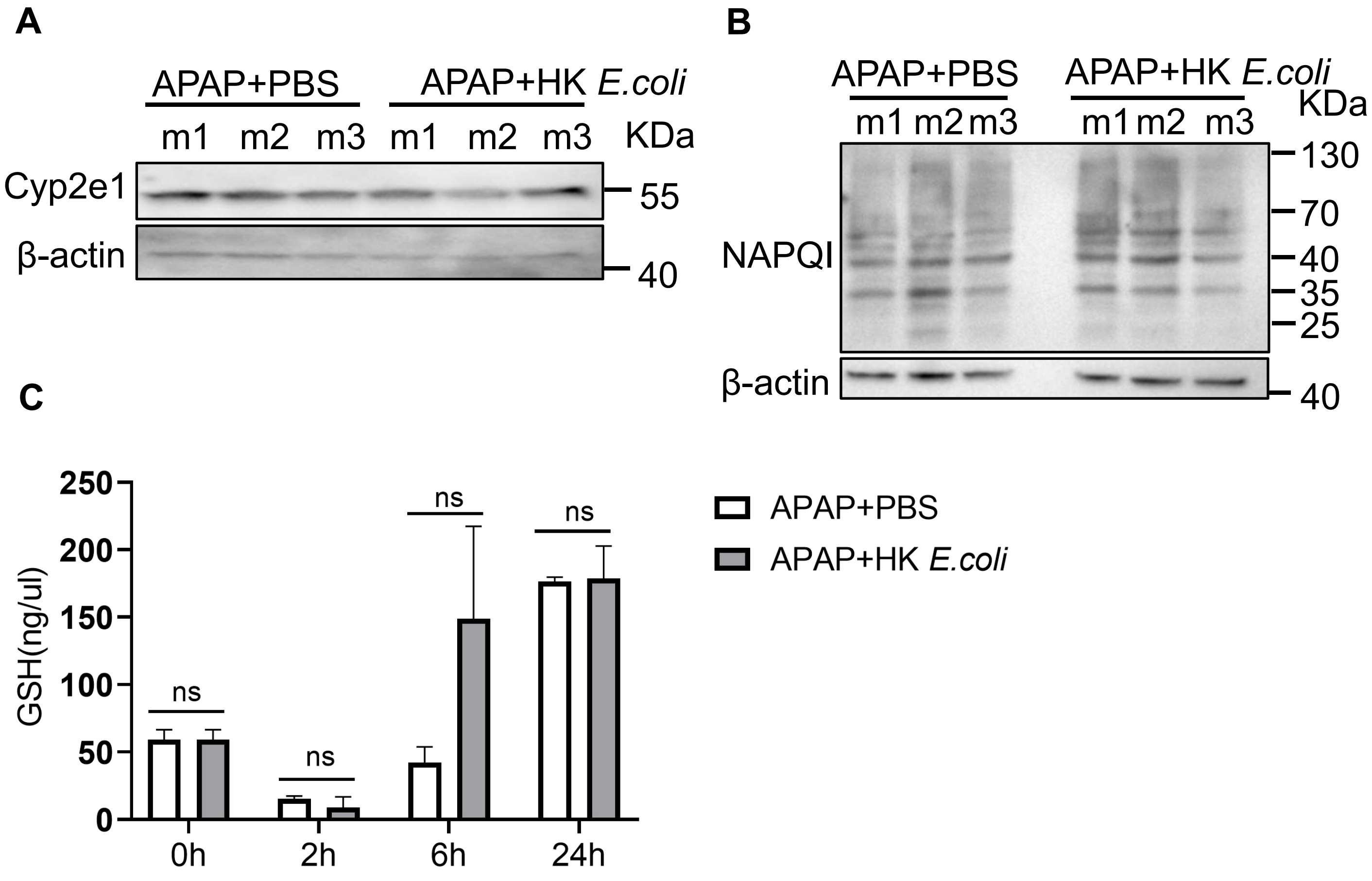


**Figure S1. HK *E. coli* does not alter APAP metabolism *in vivo*.**

**(A)** Western blot analysis of CYP2E1 protein levels in liver tissues from mice treated with APAP + PBS or APAP + HK *E. coli* at 2h post-APAP. (*n* = 3 mice/group).

**(B)** Western blot analysis of APAP adducts (NAPQI) in liver tissues from mice treated as in (A). (*n* = 3 mice/group).

**(C)** Hepatic glutathione (GSH) levels measured by HPLC at 0, 2, 6, and 24 h post-APAP in mice treated with APAP + PBS or APAP + HK *E. coli*.

Unpaired Student's t-test (C); ns, not significant ($P \geq 0.05$).

Figure S2

A

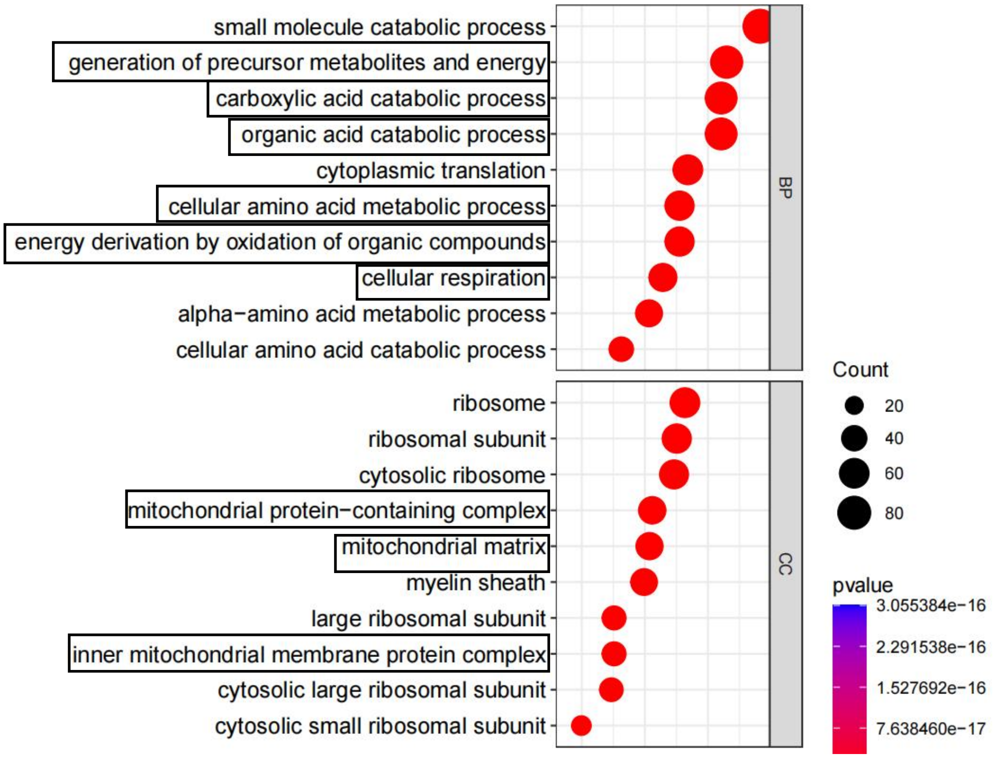


**Figure S2 | Mitochondrial functional enrichment of PGN-interacting proteins**

**(A)** Gene Ontology (GO) enrichment analysis of PGN-binding proteins identified by mass spectrometry. Bubble plot displays the top 10 significantly enriched terms in biological processes (BP, top) and cellular components (CC, down). Mitochondria-related process or components were outlined in black rectangles. Full mass spectrometry data are available in Table 1.

#### Figure S3

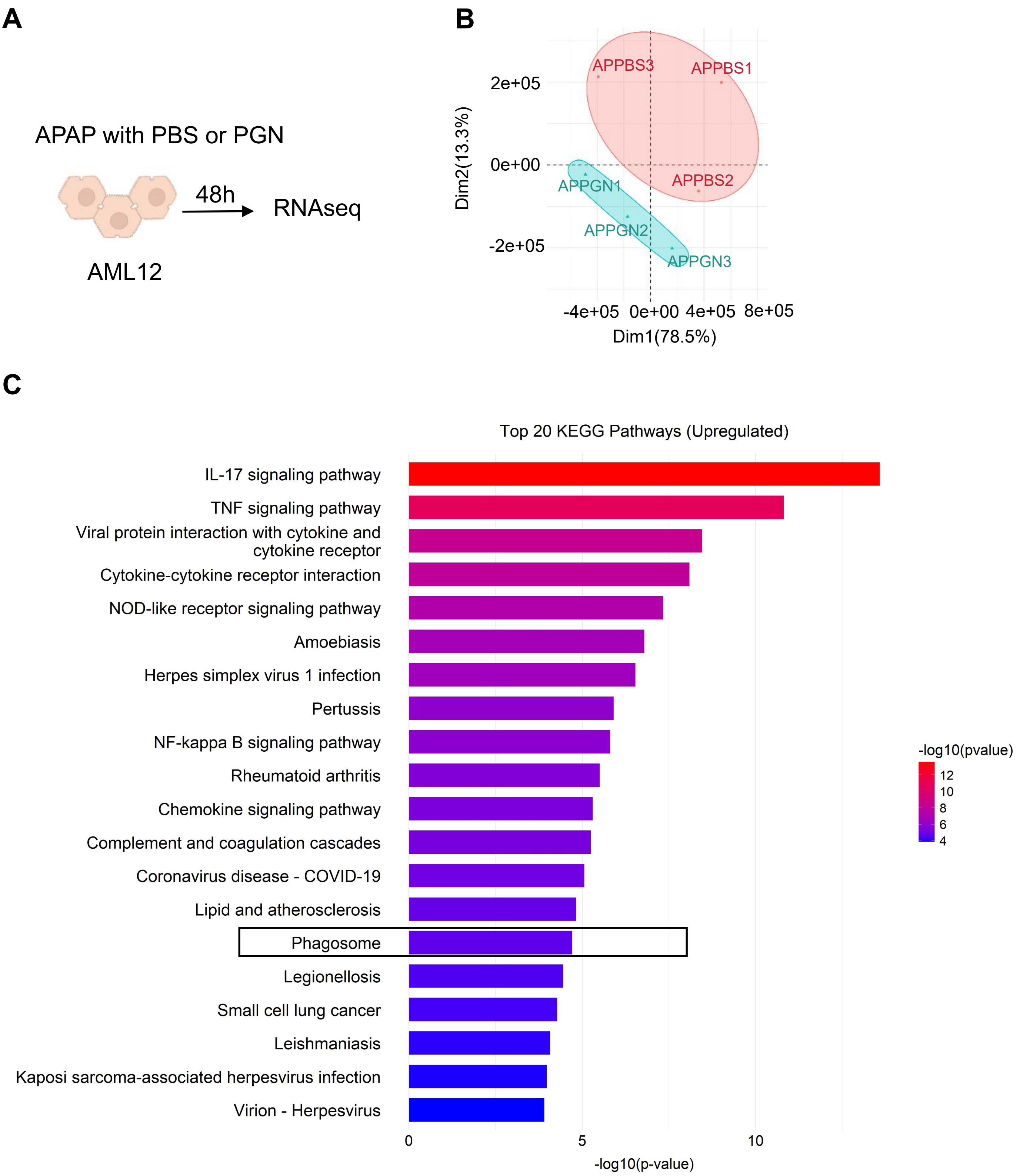

A
APAP with PBS or PGN
48h
RNAseq
AML12
B
APPBS3
APPBS1
APPBS2
APPGN1
APPGN2
APPGN3
2e+05
0e+00
-2e+05
Dim2(13.3%)
-4e+05
0e+00
4e+05
8e+05
Dim1(78.5%)
C
Top 20 KEGG Pathways (Upregulated)
IL-17 signaling pathway
TNF signaling pathway
Viral protein interaction with cytokine and cytokine receptor
Cytokine-cytokine receptor interaction
NOD-like receptor signaling pathway
Amoebiasis
Herpes simplex virus 1 infection
Pertussis
NF-kappa B signaling pathway
Rheumatoid arthritis
Chemokine signaling pathway
Complement and coagulation cascades
Coronavirus disease - COVID-19
Lipid and atherosclerosis
Phagosome
Legionellosis
Small cell lung cancer
Leishmaniasis
Kaposi sarcoma-associated herpesvirus infection
Virion - Herpesvirus
0
5
10
-log10(p-value)
-log10(pvalue)
12
10
8
6
4

**Figure S3. PGN enriches phagosome-related biological processes in transcriptomic analysis.**

**(A)** Experimental design for RNA-seq in AML12 cells treated with APAP with either PBS or PGN.

**(B)** Principal component analysis (PCA) of RNA-seq data, showing distinct transcriptional profiles between treatment groups.

**(C)** KEGG pathway enrichment analysis of upregulated differentially expressed genes (DEGs) in APAP+PGN versus APAP+PBS. The y-axis displays top 20 enriched KEGG pathways, with the phagosome pathway highlighted by a black box. The x-axis represents the GeneRatio, calculated as the number of enriched genes divided by the total number of genes in the pathway. The color scale (right) represents the adjusted *q*-value, with darker colors indicating higher statistical significance. Circle size reflects the number of DEGs mapped to each pathway. Full RNA-seq data is availble in Table 2.

### Figure S4

**A**

autophagy-related proteins

| Genes names | Peptides |
|---|---|
| TUFM | 6 |
| Ubc; Ubb; Rps27a; Uba52; GM8797; Kxd1 | 4 |
| Vdac1 | 4 |
| Rps27l; Rps27 | 2 |
| Vdac2 | 1 |
| Vdac3 | 1 |
| P62 | 1 |

**Figure S4. Autophagy-related proteins pulled down by PGN.**

**(A)** List of autophagy-related proteins identified as PGN interactors by mass spectrometry. Full mass spectrometry data are available in Table 1.

**Figure S5**

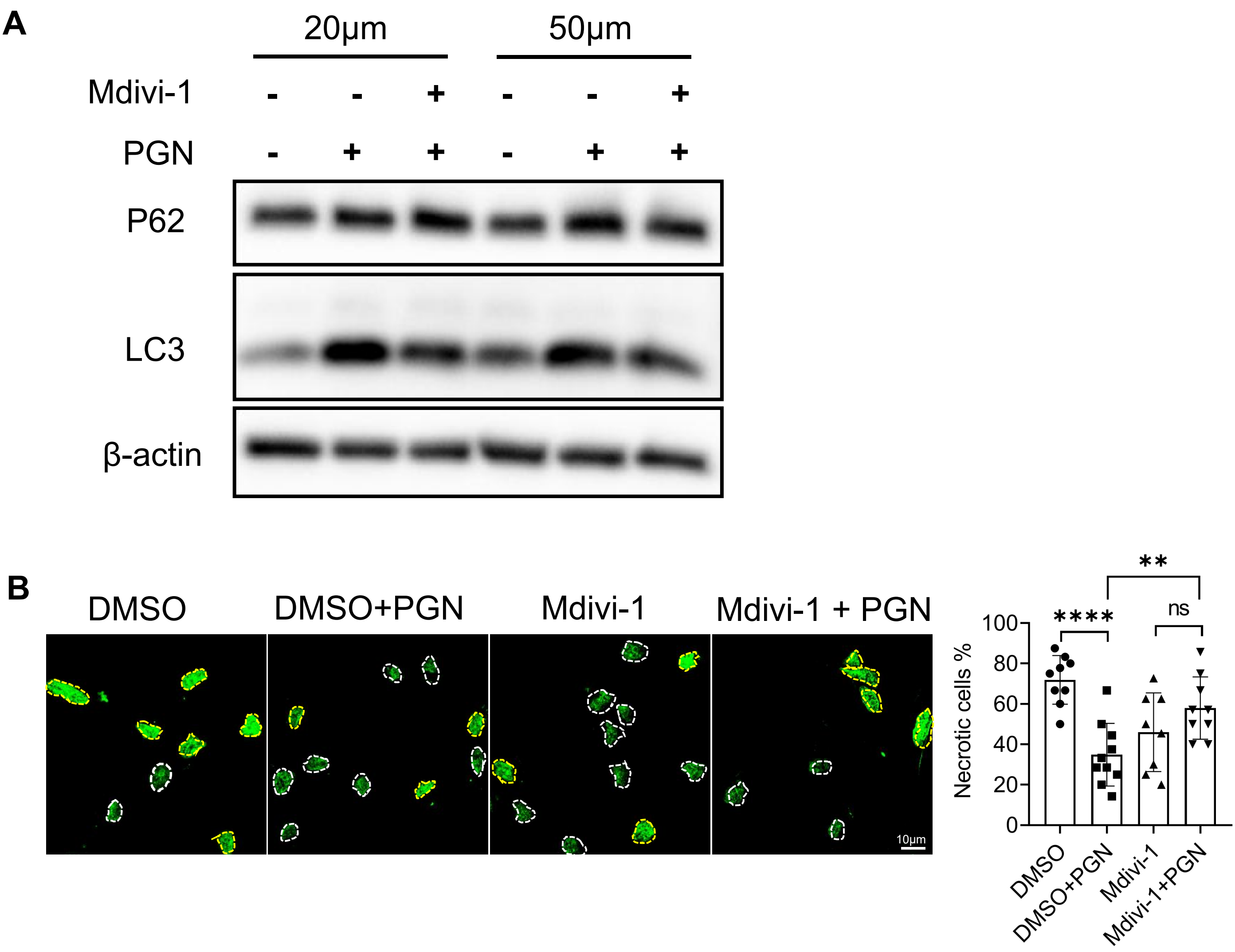


**Figure S5. PGN-mediated inhibition of cell death requires mitophagy**

**(A)** Immunoblot analysis of autophagy-related proteins in Huh7 cells treated with Mdivi-1 (20 μM or 50 μM) in the presence or absence of PGN.

**(B)** SYTOX Green staining to assess cell death in Huh7 cells treated with DMSO, DMSO + PGN, Mdivi-1, or Mdivi-1 + PGN. Left: Representative fluorescence images (yellow dashed outline: SYTOX Green-positive dead cells; white dashed outline: viable cells). Right: Quantification of necrotic cells. Data are mean ± SEM (*n* = 9 images per group). Statistical significance was determined by one-way ANOVA (ns, not significant, $P \geq 0.05$; $^{**}P < 0.01$; $^{****}P < 0.0001$).
